# Fast Principal Component Analysis of Large-Scale Genome-Wide Data

**DOI:** 10.1101/002238

**Authors:** Gad Abraham, Michael Inouye

**Affiliations:** Medical Systems Biology, Department of Pathology and Department of Microbiology & Immunology, University of Melbourne, Parkville 3010, Victoria, Australia

## Abstract

Principal component analysis (PCA) is routinely used to analyze genome-wide single-nucleotide polymorphism (SNP) data, for detecting population structure and potential outliers. However, the size of SNP datasets has increased immensely in recent years and PCA of large datasets has become a time consuming task. We have developed `flashpca`, a highly efficient PCA implementation based on randomized algorithms, which delivers identical accuracy in extracting the top principal components compared with existing tools, in substantially less time. We demonstrate the utility of `flashpca` on both HapMap3 and on a large Immunochip dataset. For the latter, `flashpca` performed PCA of 15,000 individuals up to 125 times faster than existing tools, with identical results, and PCA of 150,000 individuals using `flashpca` completed in 4 hours. The increasing size of SNP datasets will make tools such as `flashpca` essential as traditional approaches will not adequately scale. This approach will also help to scale other applications that leverage PCA or eigen-decomposition to substantially larger datasets.

## Introduction

Principal component analysis (PCA) is a widely-used tool in genomics and statistical genetics, employed to infer cryptic population structure from genome-wide data such as single nucleotide polymorphisms (SNPs) [1, 2], and/or to identify outlier individuals which may need to be removed prior to further analyses, such as genome-wide association studies (GWAS). This is based on the fact that such population structure can confound SNP-phenotype associations, resulting in some SNPs spuriously being called as associated with the phenotype (false positives). The top principal components (PCs) of SNP data have been shown to map well to geographic distances between human populations [1, 3], thus capturing the coarse-grain allelic variation between these groups.

However, traditional approaches to computing the PCA, such as those employed by the popular EIGENSOFT suite [1], are computationally expensive. For example, PCA based on the singular value decomposition (SVD) scales as *𝒪*(min (*N*^2^*p, N p*^2^)) and for eigen-decomposition it is *𝒪*(min (*N*^3^, *p*^3^)) (excluding the cost of computing the covariance matrix itself which is also *𝒪*(min (*N*^2^*p, N p*^2^))), where *N* and *p* are the number of samples and SNPs, respectively. This makes it time-consuming to perform PCA on large cohorts such as those routinely being analyzed today, involving millions of assayed or imputed SNPs and tens of thousands of individuals, with this difficulty only likely to increase in the future with the availability of even larger studies.

In recent years, research into randomized matrix algorithms has yielded alternative approaches for performing PCA and producing these top PCs, while being far more computationally tractable and maintaining high accuracy relative to the traditional “exact” algorithms [4, 5]. These algorithms are especially useful when we are interested in finding only the first few principal components (PCs) of the data, as is often the case in genomic analysis.

Here we present `flashpca`, an efficient tool for performing PCA on large genome-wide data, based on randomized algorithms. Our approach is highly efficient, allowing the user to perform PCA on large datasets (100,000 individuals or more), extracting the top principal components while achieving identical results to traditional methods.

## Results

First we used an LD-pruned HapMap3 genotype data [6] consisting of 957 human individuals across 11 populations assayed for 14,389 SNPs (Materials). We compared `flashpca` with `smartpca` from EIGENSOFT v4.2^1^ and shellfish^2^. In addition, we included the `R` 3.0.2 `prcomp` function which is based on SVD rather than eigen-decomposition, after replacing its original standardization with the one used by `smartpca` (Equation 4). The analysis of HapMap3 data revealed the expected ancestry via the first two PCs (Figure 1a), with individuals of east Asian origin (CHB, CHD, JPT) clustered in the bottom right-hand side corner, those of European origin (TSI, CEU) in the top, and those of African origin in the bottom left-hand side corner (ASW, LWK, YRI, and MKK). All methods showed close to perfect agreement on the top PC (Figure 1b), with an absolute correlation of 1.00 between the 1st PC of each pair of methods (the sign of the eigenvectors is arbitrary hence the correlation may be negative as well). The next nine PCs showed close to perfect agreement as well (see Supplementary File 1 and the online documentation for results for PCs 1–10).

**Figure 1:**
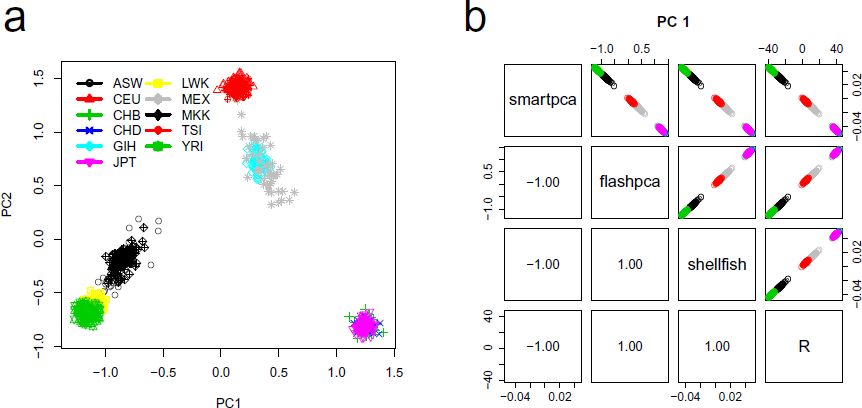
(a) The first two principal components from analyzing the HapMap3 dataset. (b) Scatter plots showing near-perfect absolute Pearson correlation (lower left-hand corner) between the 1st PCs estimated by `smartpa`, `flashpca`, `shellfish`, and `R`’s `prcomp` (using the standardization from Equation 4). Note that since eigenvectors are only defined up to sign, the correlations may be negative as well as positive. In addition, the scale of the PCs may differ between the methods, however, this has no bearing on the interpretation of the PCs.

Next, we analyzed an LD-pruned celiac disease Immunochip dataset consisting of 16,002 individuals and 43,049 SNPs after thinning by LD [7] (Materials). We then randomly sampled subsets of the original dataset with increasing size (*N* = 500, 1000, 2500, 5000, 7500, 10,000, 15,000), and recorded wall time for `flashpca`, `smartpca`, and `shellfish` performing PCA on these subsets. We also examined larger setups (*N* = 50,000, 100,000, and 150,000) by duplicating the original dataset several times as required.

Due to the substantial time required by `shellfish` and `smartpca` to complete the largest runs, we only ran `flashpca` on the larger datasets (*N ≥* 50,000) (we attempted to run `shellfish` on the *N* = 50,000 dataset but it did not complete due to running out of memory, and we stopped `smartpca` after 100,000sec).

Each experiment was repeated three times. All programs used multi-threaded mode with 8 threads. All experiments were run on a machine with 4*×*10-core Intel Xeon E7-4850 CPU @ 2.00GHz with 512GiB RAM running 64-bit Ubuntu Linux 12.04. Note that wall time here is defined as time from program start to successful exit, inclusive of any loading of data, scaling, computation of the covariance matrix, eigen-decomposition, and so on, however, the majority of the run time is taken by computing the covariance and the eigen-decomposition. `smartpca` was run without excluding potential population outliers.

Figure 2 shows that `flashpca` was substantially faster than either `smartpca` or `shellfish`: for analysis of 15,000 samples, `flashpca` took an average of 8 minutes, whereas `smartpca` required an average of almost 17h (*×*125 slower). Examining the large datasets, `flashpca` was able to analyze a dataset of *N* = 150,000 in *∼*4h, which would not be sufficient time for `smartpca` to complete a PCA on 10,000 samples, as 6.5h were required for that. While `shellfish` was also substantially faster than `smartpca`, it was still considerably slower than `flashpca` when the number of individuals was large, with a run time of *∼* 1h for *N* = 15,000 (and did not complete for subsets with *N ≥* 50,000).

**Figure 2:**
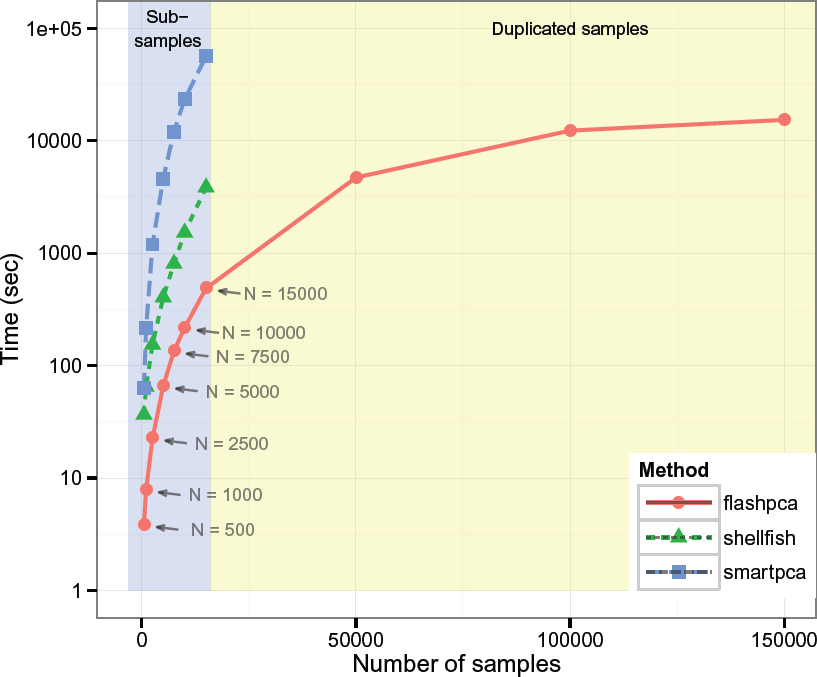
Total wall time (seconds) for `flashpca` versus EIGENSOFT’s `smartpca` and `shellfish` on increasing subsets of the celiac disease dataset, employing multi-threaded mode (8 threads), using 43,049 SNPs. `shellfish` did not complete PCA for the *N ≥* 50,000 subsets, and `smartpca` was stopped after 100,000sec. The results shown are averages over 3 runs. Results for *N ≤* 15,000 are based on subsamples of the original dataset *N* = 16,002 (light blue background), whereas results for *N ≥* 50,000 are based on duplicating the original samples (light yellow background).

Importantly, the time taken by `flashpca` to compute PCA on *N* = 150,000 did not differ much from the time taken on the *N* = 100,000 subset; this is because `flashpca` automatically transposes the data when *N > p*, as the PCA can be computed on the original data or its transpose with only minor modifications to the algorithm; computing the *p × p* covariance matrix and its eigen-decomposition has lower computational complexity than using the *N × N* covariance matrix, for the same values of *N* and *p*, hence when *N > p* the main computational cost will not grow substantially with *N*.

## Discussion

Principal component analysis is an important tool in genomics for discovery of population structure or other latent structure in the data, such as batch effects. Early approaches such as `smartpca` from EIGENSOFT have proven useful for this goal and have been widely used for analysis of SNP datasets. However, many current datasets assay tens of thousands of individuals, making traditional approaches extremely time consuming. In contrast, our approach, `flashpca`, is based on careful combination of randomized algorithms for PCA together with parallelization, and allows the analyst to easily perform PCA on large datasets consisting of many thousands of individuals in a matter of minutes to hours. Despite relying on an approximation strategy, this approach suffers from essentially no loss in accuracy for the top eigenvalues/eigenvectors compared with traditional approaches. One practical limitation of the current implementation of `flashpca` is its memory requirements for large datasets: using 15,000 individuals with 43K SNPs requires *∼*14GiB RAM, and 150,000 requires *∼*145GiB. Future work will involve reducing these memory requirements without incurring a substantial performance penalty. Randomized PCA also provides the potential to de-correlate samples (“whitening”), thus essentially removing the effects of population from data prior to further downstream association analysis [8]; this will be examined in future work. It has been shown that standard PCA may be an inconsistent estimator of the true principal components in certain high dimensional settings [9], and there may be benefit from utilizing other approaches such as sparse (*ℓ*_1_-penalized) PCA [10, 11]; however, standard PCA remains widely used in practice and `flashpca` provides an effective way to perform such routine analyses highly efficiently.

More generally, the approach behind `flashpca` could prove useful for accelerating other methods that depend on performing a large number of eigen-decompositions across many samples, such as varLD [12] which assesses local differences in SNP LD between populations or FaST-LMM which implements linear mixed models of SNP data [13].

## Materials and Methods

### Ethics

All subjects included in the celiac disease dataset provided written and informed consent. For details, see the original publication [7].

### Datasets

#### HapMap3

The HapMap phase 3 dataset consists of 1184 human individuals across 11 populations (ASW: African ancestry in Southwest USA; CEU: Utah residents with Northern and Western European ancestry from the CEPH collection; CHB: Han Chinese in Beijing, China; CHD: Chinese in Metropolitan Denver, Colorado; GIH: Gujarati Indians in Houston, Texas; JPT: Japanese in Tokyo, Japan; LWK: Luhya in Webuye, Kenya; MEX: Mexican ancestry in Los Angeles, California; MKK: Maasai in Kinyawa, Kenya; TSI: Toscani in Italia; YRI: Yoruba in Ibadan, Nigeria) assayed for 1,440,616 SNPs [6]. We performed QC on the data, including removal of SNPs with MAF*<* 1%, missingness *>* 1%, and deviation from Hardy-Weinberg equilibrium *P <* 5 *×* 10^−6^. We removed non-founders and individuals with genotyping missingness *>* 1%, leaving 957 individuals. Next, we removed several regions of high LD and/or known inversions (chr5: 44Mb–51.5Mb, chr6: 25Mb–33.5Mb, chr8: 8Mb–12Mb, chr11: 45Mb–57Mb) [14]. Finally, we used PLINK [15] `--indep-pairwise` 1000 10 0.02 to thin the SNPs by LD (*r*^2^ *<* 0.02), leaving 14,389 SNPs.

##### Celiac Disease Immunochip

The celiac disease Immunochip dataset [7] consists of 16,002 case/control individuals of British descent, assayed for 139,553 SNPs. The QC has been previously described [7]. In addition, we removed SNPs with MAF *<* 0.5% and non-autosomal SNPs, leaving 115,746 SNPs. Next, we removed the same four regions as for the HapMap3 data, and finally, we thinned the SNPs by LD with PLINK `--indep-pairwise` 5010 0.5, leaving 43,049 SNPs.

### Principal Component Analysis

We represent the genotypes as an *N × p* matrix **X**, where the *N* samples index the rows and the *p* SNPs index the columns. We denote the transpose of the matrix as **X**^*T*^. For the following, we assume that the matrix **X** has been centered so that the mean of each column *j* is zero (see below for variants on this).

PCA relies on finding the eigenvectors of the *N × N* covariance matrix **Σ** = **XX**^*T*^ (for notational clarity we will ignore the scaling 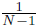 factor which can be accounted for later). This decomposition is performed using either the singular value decomposition (SVD) of the matrix **X** or the eigen-decomposition of the covariance matrix itself.

The SVD of **X** is

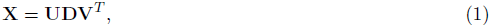

where **U** an is an *N × k* matrix (**U**^*T*^ **U** = **I**) the columns of which are the eigenvectors of **XX**^*T*^, **D** is a *k × k* diagonal matrix of singular values (square root of the eigenvalues of **XX**^*T*^ and **X**^*T*^**X**), and **V** is a *p × k* matrix (**V**^*T*^ **V** = **I**) of the eigenvectors of **X**^*T*^ **X**, where *k* is the matrix rank (this SVD is also called the “economy SVD”). Note that SVD does not require the covariance matrix **Σ** to be computed explicitly.

In the eigen-decomposition approach, the covariance matrix **Σ** is first explicitly computed, then the eigen-decomposition is performed such that

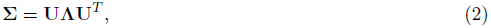

where diag(**Λ**) = *λ*_1_,…, *λ*_*k*_ = diag(**D**^2^) are the eigenvalues and **U** is the matrix of eigenvectors as before.

The principal components (PCs) **P** of the data are given by the projection of the data onto the eigenvectors

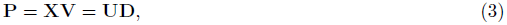

where we usually truncate the matrix **P** to have as many columns as required for any down-stream analysis (say, 10).

Note that some tools, such as `smartpca` and `shellfish`, output the eigenvectors **U** as the principal components without weighting by the singular values **D**, leading to different scales for the PCs. In addition, since the (empirical) covariance is typically scaled by a factor of 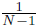, then in order to maintain the interpretation of the singular values **D** as the square-root of the eigen-values of the scaled covariance 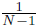**XX**^*T*^, the singular values must be scaled by a factor of 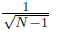 as well (as implemented in `R`’s `prcomp`).

Note, however, that these scale differences have no effect on the interpretation of the principal components for ascertaining or correcting for potential population structure in data.

In traditional PCA, such as that implemented in `R`’s `prcomp`, prior to running the SVD/eigen-decomposition itself, the matrix **X** is first mean-centered by subtracting the per-column (SNP) average from each column. In contrast, `smartpca` [1], first centers the data, then divides by a quantity proportional to the standard deviation of a binomial random variable with success probability *p*_*j*_

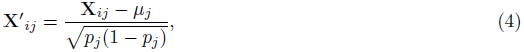

where *p_j_* = *µ_j_*/2 and 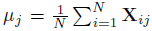. To maintain compatibility with `smartpca`, `flashpca` employs the same standardization method by default (other scaling methods are available as well).

#### Algorithm 1

Pseudocode for the eigen-decomposition variant of the fast PCA, based on the randomized algorithm of [5] for the case where *N < p*. standardize(*·*) is the standardization in Equation 4. randn(*m, n*) is a function generating an *m × n* iid multivariate normal matrix, *e* is the user-defined number of extra dimensions, qr(*·*) is the QR decomposition, normalize(*·*) is a function that divides each column *j* by its *ℓ*_2_ norm 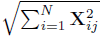, eigen(*·*) is the eigen-decomposition producing the *d*-top eigenvectors 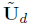 and vector of *d*-top eigenvalues ***λ***_*d*_. **P**_*d*_ is the matrix of *d* principal components.

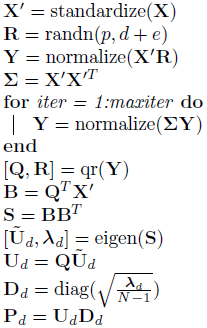

### Fast Principal Component Analysis

Performing PCA on large matrices (with both *N* and *p* large), is time consuming with traditional approaches. To enable fast PCA, we employ an algorithm based on a randomized PCA approach [5]. Briefly, randomized PCA relies on first constructing a relatively small matrix that captures the top eigenvalues and eigenvectors of the original data, with high probability. Next, standard SVD or eigen-decomposition is performed on this reduced matrix, producing near identical results to what would have been achieved using a full analysis of the original data. Since in most genomic applications we are interested only in a few of the top eigenvectors of the data (typically 10), this allows reduction of the data to a substantially smaller matrix, and the computational cost of decomposing this matrix is negligible (see Algorithm 1). (Note, however, that this method is general and is just as useful for extracting a much larger number of PCs).

Focusing on the eigen-decomposition approach (Equation 2), the two main computational bottlenecks are (i) computing the *N × N* covariance matrix **Σ** and (ii) when *N* is large, the eigen-decomposition step itself. In our fast PCA approach, the first bottleneck cannot be avoided but can be mitigated through parallel computation (see Implementation). The second bottleneck is circumvented via the randomized approach, by constructing a matrix **B** of size (*d* + *e*) *× p*, from which the small (*d* + *e*) *×* (*d* + *e*) matrix **BB**^*T*^ is formed and used for the eigen-decomposition, where *d* is the number of required eigenvectors (say 10), and *e* is the number of auxiliary dimensions used to increase accuracy which can be discarded later. We have found that a total dimension of *d* + *e* = 200 is more than sufficient for producing good results in the first 10 PCs; hence the eigen-decomposition need only be performed on 200 *×* 200 matrix while producing near identical results to a full PCA on the original data.

Another computational shortcut for the case where *N > p*, applying equally to PCA and to randomized PCA, is simply transposing the data **X**, then standardizing the rows instead of the columns. An identical PCA algorithm is then run on the transposed data, with the only difference being that the estimated matrix **U**_*d*_ will now contain the top-*d* eigenvectors of **X**′^*T*^ **X**′ instead of **X**′**X**′^*T*^, and hence the final *d* principal components will be **P**_*d*_ = **X**′**U**_*d*_. This procedure makes it possible to analyze large datasets with *N ≫ p* at a cost not much greater than when *N* = *p*.

While the SVD approach is generally recommended for reasons of better numerical stability and speed, we have found that when *N* and *p* are in the thousands, the above eigen-decomposition approach is substantially faster since the matrix multiplication is trivially parallelizable (and is parallelized in practice in `flashpca`), allowing the decomposition to be performed on the small matrix **S** = **BB**^*T*^ which is of size (*d* + *e*) *×* (*d* + *e*) rather than a more expensive non-parallelized SVD of the matrix **B**, which is an *N ×* (*d* + *e*) matrix, with no discernible effect on accuracy of the top principal components; however, both methods are implemented in `flashpca` (yet another possibility is to perform SVD on **S**, but this is not implemented yet).

### Implementation

`flashpca` is implemented in C++ and relies on Eigen [16], a C++ header-only library of numerical linear algebra algorithms, which allows for native parallelization of certain computations through OpenMP threads when multiple CPU cores are available. `flashpca` natively reads PLINK [15] SNP-major BED files, avoiding the need to convert these files to other formats. `flashpca` is licensed under the GNU Public License (GPL) v3; source code and documentation are available at https://github.com/gabraham/flashpca.

Prior to PCA, thinning of the SNPs by LD and removal of regions with known high LD or other artefacts such as inversions have been recommended [1, 14], as high correlations between the SNPs can distort the resulting eigenvectors. For this purpose we recommend using PLINK v2 3 which is substantially faster than PLINK v1.07. Reducing a dataset to *∼* 10,000–50,000 SNPs is usually sufficient to achieve an accurate PCA, and can be done using `--indep-pairwise`.

## Acknowledgments

We acknowledge support and funding from NHMRC grant no. 1062227. MI was supported by an NHMRC Early Career Fellowship (no. 637400). We thank the chief investigators of celiac disease Immunochip study (David van Heel and Cisca Wijmenga) for kindly providing the data.

## Supporting Information

**File S1:** Concordance in principal components 1–10 between `smartpca`, `flashpca`, `shellfish`, and `R`’s `prcomp` on the HapMap3 dataset.

## Footnotes

1 http://www.hsph.harvard.edu/alkes-price/software/

2 http://www.stats.ox.ac.uk/∼davison/software/shellfish/shellfish.php

3 https://www.cog-genomics.org/plink2

